# NEMETEX: a Python software for the visualisation of the network of metabolic exchanges

**DOI:** 10.1101/2022.10.19.492777

**Authors:** Michela Palamin, Alice Frisinghelli, Elisabetta Offer, Guido Zampieri, Arianna Basile, Stefano Campanaro

## Abstract

**Motivation:** Microorganisms have a pivotal role in ecology and human health and form complex networks where different species can interact and exchange a range of different compounds. Flux balance analysis can offer an insight into the production and the absorption of these metabolic compounds, but frequently results are difficult to visualise and interpret. Additionally, a clear understanding of the roles of microbial species in the community requires the integration of different information sources, including relative abundance, taxonomy and compounds exchange rate.

**Results:** To fill-in this gap the command-line tool NEMETEX (NEtwork for METabolic Exchanges) was developed to provide a graphical representation of the metabolites exchanged, joined with interactive visualisation of numerical data. This approach can undoubtedly represent an easy way to investigate high-throughput results obtained from metagenomics and flux balance analysis, providing a more direct interpretation of the data.

**Availability and implementation:** This program, accessory utilities, and their documentation are freely available at https://github.com/palakela/NEMETEX

## 1 Introduction

Microbial communities are composed of groups of microorganisms sharing the same living space and interacting with each other. In microbiomes, different species can compete for resources and exchange metabolites, two processes representing the main drivers of microbial community assembly (Ponomarova & Patil, 2015). The interplays between microorganisms are extremely variable and can range from competitive, to commensal and mutualistic interactions (Basile *et al*, 2020). A complete understanding of these phenomena is limited by the difficulty in performing direct experimental measurement of carbon or nitrogen fluxes, however, the advancement of metagenomic techniques combined with metabolic Flux Balance Analysis (FBA) (Orth *et al*, 2010) allowed the investigation of metabolic fluxes within communities at an unprecedented level of resolution (Machado *et al*, 2021; Diener *et al*, 2020). The recent improvement of methods focused on recovery of Metagenome-Assembled Genomes (MAGs) from shotgun sequencing determined a massive recovery of uncharacterized microbial species, most of them having a yet unclarified role (Hugenholtz & Tyson, 2008; Forster *et al*, 2019). Additionally, recently developed tools such as CarveMe and gapseq (Machado *et al*, 2018; Zimmermann *et al*, 2021) allow a fast and easy generation of genome-scale metabolic models, and can provide mechanistic bases for the prediction of metabolites exchange among species. The use of these models in mathematical modelling frameworks can also simulate metabolic interactions (Diener *et al*, 2020; Bauer *et al*, 2017). However, the output files obtained from these analyses are of difficult interpretation, and therefore the possibility of obtaining a graphical visualisation of the results can help in understanding the different functional roles of the microbial species. Approaches for the graphical visualisation of metabolites exchanged in complex communities accounting for more than two microbial species, are already existing (Bauer *et al*, 2017; Diener *et al*, 2020). However, these software require the import of results from flux balance simulations in a separated environment, thus requesting programming knowledge. Furthermore, these systems are not designed for metagenomic data, and, as a consequence, they don’t include information regarding microbial abundances and taxonomic assignment, which are compulsory for a meaningful biological interpretation of the results. To fill this gap, we have developed NEMETEX (NEtwork for METabolic EXchanges), a Python script to graphically visualise the metabolites exchanged in a microbiome obtained through FBA simulations, integrating features of the microbial community like taxonomy and species abundance.

## 2 Results

NEMETEX is a tool implemented in Python 3, and developed in order to obtain a visual representation of the potential cross-feedings between community members. The source code is available in the dedicated GitHub repository and for its functioning the libraries Numpy v1.15.1, Pandas v0.23.4, NetworkX v2.1., PyVis v0.1.9, argparse v2.0.1, os are needed.

As depicted in the workflow diagram illustrating the pipeline (Fig. 1A), the script requires three input files reporting (i) the compounds exchanged, (ii) the species abundance and (iii) the taxonomic assignment for the species.

i. Metabolites exchange rate is imported from the output file (.tsv format) generated by SMETANA software (v1.0.0) (Species METabolic interaction ANAlysis) (Zelezniak *et al*, 2015). The “receiver” and the “donor” species are identified along with the exchange rate (smetana value) of each compound from the tabular file.
ii. Relative abundance of each species is imported as a tabular file generated using checkM (v1.0.12) (Parks *et al*, 2015).
iii. The species taxonomy is defined from GTDB-Tk (v1.3.0) software (Chaumeil *et al*, 2019) and converted to the NCBI taxonomy using the script gtdb_to_ncbi_majority_vote.py.

**Figure 1.**
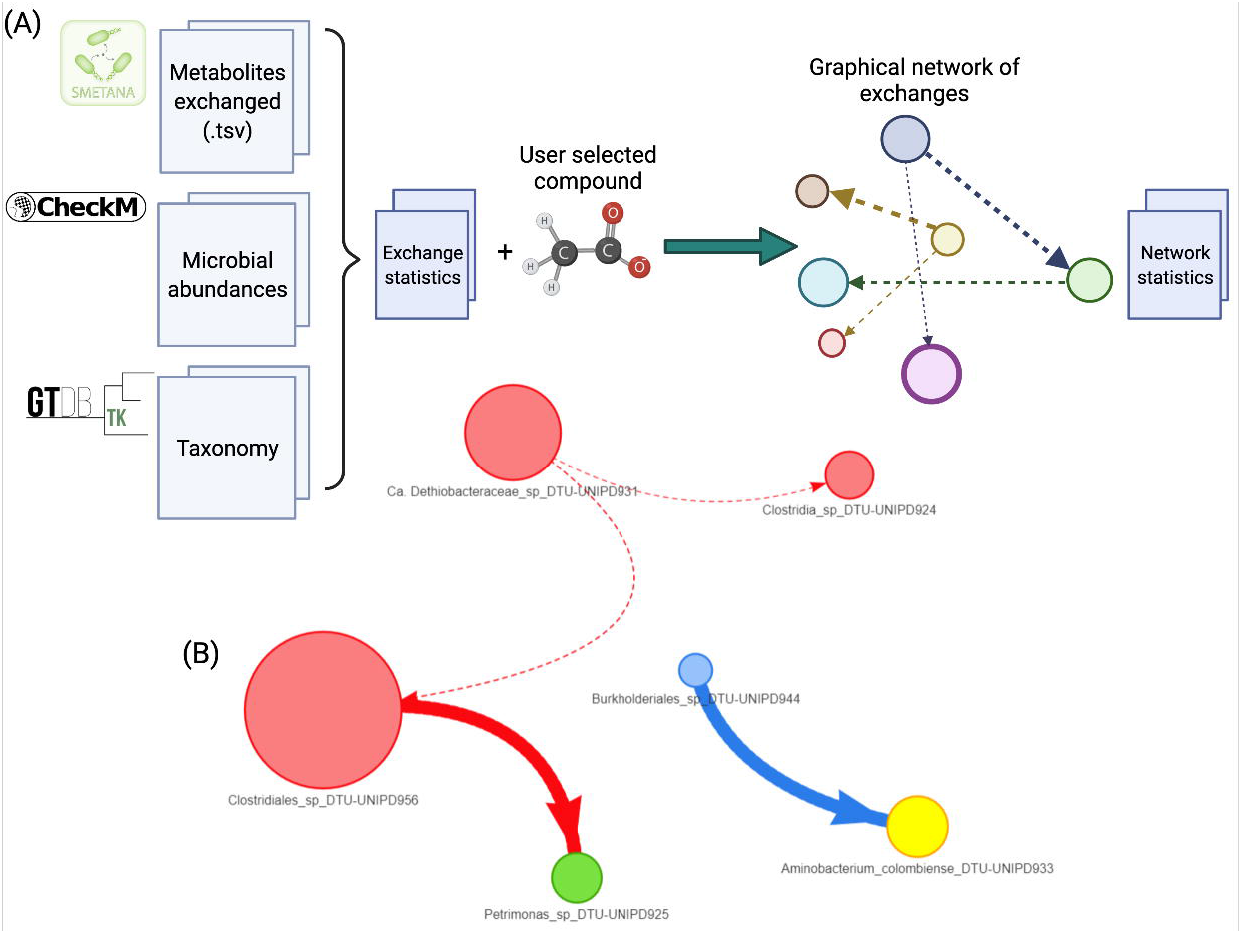
(A) NEMETEX workflow. (B) Butyrate exchanges in the microbiome under investigation. Colour and size of each node refer respectively to taxonomic assignment at phylum level and species relative abundance. Arrows reflect directionality of metabolite exchange, whereas thickness indicates exchange probability.

The script provides as output not only the network in html format, but also three global summary files reporting for all compounds the number of exchanges for each compound, the donor and receiver species. A preliminary inspection of these latter files can be helpful in order to select what are the compounds the user would like to investigate more in detail with the graphical representation.

Finally, the tool requires the compound of interest (extended name, biggID, or ModelSEED ID) as input to visualise how it is exchanged within the community. The image reporting the network is generated together with two files describing the behaviour of the species (donor-receiver) in the exchange of the compound of interest and reporting the exchanges information. It is also possible to provide the tool with a text file with the list of compounds, one per line. In the latter case, NEMETEX will generate the three output files for each compound in separate directories.

## 3 Applications

The main output provided by NEMETEX consists of an HTML file reporting an interactive network that allows to visualise in an easy way the metabolic exchanges of the compounds of interest. In the network, the nodes represent the microbial species, with edges showing the directional exchanges between them. The HTML file generated can be used for the dynamic visualisation of the network (Supplementary file I). The graphical representation is tailored to help the user in the identification of the network, and simplify data interpretation. The diameter of each node is proportional to the species relative abundance for an easier identification of dominant species in the community. In the final rendering, the nodes are colored according to the taxonomic assignment at phylum level to facilitate the investigation on specific taxonomic groups. Arrows thickness refers instead to the probability of exchange as calculated by SMETANA (Zelezniak *et al*, 2015), and the relative value can be directly displayed by moving the mouse cursor over the edge. Additional information can be visualised by moving the mouse cursor over the nodes, including for example the taxonomic assignment of the species at phylum level, their relative abundance, and the species receiving the compound under investigation. The user will be further supported in the interpretation of the results thanks to the supplementary output tabular files reporting some basic statistics calculated from SMETANA output.

The tool was tested with results obtained from a genome-centric metagenomic analysis performed on samples collected from an anaerobic biogas reactor (De Bernardini *et al*, 2022). As an illustrative example of the NEMETEX output format, analysis of the butyrate exchanges within the selected microbial community, is shown in figure 1B. In this example, there are four main phyla involved, *Firmicutes* is colored in red, *Proteobacteria* in blue, *Synergistetes* in yellow, and *Bacteroidetes* in green. Five of the reconstructed genomes cannot be assigned at species level by GTDB-Tk, while only *Aminobacterium colombiense* was clearly assigned to the species level. The network has to be interpreted as follows: three donor species were identified (Clostridiales sp. DTU-UNIPD956, Candidatus Dethiobacteraceae sp. DTU-UNIPD931 and Burkholderiales sp. DTU-UNIPD944), and three receivers (*Aminobacterium colombiense* DTU-UNIPD933, Petrimonas sp. DTU-UNIPD925 and Clostridiales sp. DTU-UNIPD956). The dominant species represented in the microbial community (Clostridiales sp. DTU-UNIPD956), with a relative abundance of 0.85%, can be easily identified from the circle diameter, as well as the main exchanges of the compounds which are evidenced by the edges thickness.

## 4 Limitations

NEMETEX has been developed considering the typical structure of the output files obtained from a genome-centric metagenomic experiment; these files include MAGs with known relative abundance calculated from alignment of the shotgun sequences on the reference assembly, and with taxonomy determined using reference genes. However, the tool can be easily adapted to the simulations performed with known species. The tool is strictly linked to the output files describing metabolites exchange and obtained using SMETANA (Zelezniak *et al*, 2015), to the relative abundance of species obtained using checkM (Parks *et al*, 2015), and to the taxonomic assignment obtained using GTDB-Tk. The tool can certainly be adapted to import output files obtained from other tools such as BAT (von Meijenfeldt *et al*, 2019) or coverM (https://github.com/wwood/CoverM), and authors are planning to include these options in future releases of NEMETEX. Additionally, if the desired compound is exchanged by a high number of species, the generated network can become very complex. In this case, the network representation encounters its limit. However, the reported statistics can give an insight on the pivotal donors and receivers.

## 5 Conclusions

NEMETEX enables a meaningful visualisation of metabolic exchanges within microbial communities and provides statistics of the FBA results. The tool is particularly suited for results obtained from SMETANA simulations. Unlike previously developed approaches, NEMETEX integrates information of metagenomic data composition and taxonomy in the network. This feature will simplify the identification of the most relevant metabolites exchange among species, and will greatly improve the biological interpretation of the results.

## 7 CRediT authorship contribution statement

**Michela Palamin:** Methodology, Software, Writing - Original Draft, Writing - Review & Editing. **Alice Frisinghelli:** Validation, Writing - Original Draft, Writing - Review & Editing. **Elisabetta Offer:** Validation, Investigation, Data curation, Writing - Review & Editing, Visualization. **Guido Zampieri:** Writing - Review & Editing. **Arianna Basile:** Conceptualization, Supervision, Methodology, Software, Visualization, Writing - Original Draft, Writing - Review & Editing. **Stefano Campanaro:** Conceptualization, Supervision, Project administration, Funding acquisition, Writing - Original Draft, Writing - Review & Editing.

## References

Basile A, Campanaro S, Kovalovszki A, Zampieri G, Rossi A, Angelidaki I, Valle G & Treu L (2020) Revealing metabolic mechanisms of interaction in the anaerobic digestion microbiome by flux balance analysis. Metab Eng 62: 138–149

Bauer E, Zimmermann J, Baldini F, Thiele I & Kaleta C (2017) BacArena: Individual-based metabolic modeling of heterogeneous microbes in complex communities. PLoS Comput Biol 13: e1005544

Chaumeil P-A, Mussig AJ, Hugenholtz P & Parks DH (2019) GTDB-Tk: a toolkit to classify genomes with the Genome Taxonomy Database. Bioinforma Oxf Engl:btz848

De Bernardini N, Basile A, Zampieri G, Kovalovszki A, De Diego Diaz B, Offer E, Wongfaed N, Angelidaki I, Kougias PG, Campanaro S & Treu L (2022) Integrating metagenomic binning with flux balance analysis to unravel syntrophies in anaerobic CO2 methanation. Microbiome 10, no. 1: 1–18.

Diener C, Gibbons SM & Resendis-Antonio O (2020) MICOM: Metagenome-Scale Modeling To Infer Metabolic Interactions in the Gut Microbiota. mSystems 5: e00606–19

Forster SC, Kumar N, Anonye BO, Almeida A, Viciani E, Stares MD, Dunn M, Mkandawire TT, Zhu A, Shao Y, et al (2019) A human gut bacterial genome and culture collection for improved metagenomic analyses. Nat Biotechnol 37: 186–192

Hugenholtz P & Tyson GW (2008) Metagenomics. Nature 455: 481–483

Machado D, Andrejev S, Tramontano M & Patil KR (2018) Fast automated reconstruction of genome-scale metabolic models for microbial species and communities. Nucleic Acids Res 46: 7542–7553

Machado D, Maistrenko OM, Andrejev S, Kim Y, Bork P, Patil KR & Patil KR (2021) Polarization of microbial communities between competitive and cooperative metabolism. Nat Ecol Evol 5: 195–203

von Meijenfeldt FAB, Arkhipova K, Cambuy DD, Coutinho FH & Dutilh BE (2019) Robust taxonomic classification of uncharted microbial sequences and bins with CAT and BAT. Genome Biol 20: 217

Orth JD, Thiele I & Palsson BØ (2010) What is flux balance analysis? Nat Biotechnol 28: 245–248

Parks DH, Imelfort M, Skennerton CT, Hugenholtz P & Tyson GW (2015) CheckM: assessing the quality of microbial genomes recovered from isolates, single cells, and metagenomes. Genome Res 25: 1043–1055

Ponomarova O & Patil KR (2015) Metabolic interactions in microbial communities: untangling the Gordian knot. Curr Opin Microbiol 27: 37–44

Zelezniak A, Andrejev S, Ponomarova O, Mende DR, Bork P & Patil KR (2015) Metabolic dependencies drive species co-occurrence in diverse microbial communities. Proc Natl Acad Sci U S A 112: 6449–6454

Zimmermann J, Kaleta C & Waschina S (2021) gapseq: informed prediction of bacterial metabolic pathways and reconstruction of accurate metabolic models. Genome Biol 22: 81

